# Accurate viral genome reconstruction and host assignment with proximity-ligation sequencing

**DOI:** 10.1101/2021.06.14.448389

**Authors:** Gherman Uritskiy, Maximillian Press, Christine Sun, Guillermo Domínguez Huerta, Ahmed A. Zayed, Andrew Wiser, Jonas Grove, Benjamin Auch, Stephen M. Eacker, Shawn Sullivan, Derek M. Bickhart, Timothy P. L. Smith, Matthew B. Sullivan, Ivan Liachko

## Abstract

Viruses play crucial roles in the ecology of microbial communities, yet they remain relatively understudied in their native environments. Despite many advancements in high-throughput whole-genome sequencing (WGS), sequence assembly, and annotation of viruses, the reconstruction of full-length viral genomes directly from metagenomic sequencing is possible only for the most abundant phages and requires long-read sequencing technologies. Additionally, the prediction of their cellular hosts remains difficult from conventional metagenomic sequencing alone. To address these gaps in the field and to accelerate the study of viruses directly in their native microbiomes, we developed an end-to-end bioinformatics platform for viral genome reconstruction and host attribution from metagenomic data using proximity-ligation sequencing (i.e., Hi-C). We demonstrate the capabilities of the platform by recovering and characterizing the metavirome of a variety of metagenomes, including a fecal microbiome that has also been sequenced with accurate long reads, allowing for the assessment and benchmarking of the new methods. The platform can accurately extract numerous near-complete viral genomes even from highly fragmented short-read assemblies and can reliably predict their cellular hosts with minimal false positives. To our knowledge, this is the first software for performing these tasks. Being significantly cheaper than long-read sequencing of comparable depth, the incorporation of proximity-ligation sequencing in microbiome research shows promise to greatly accelerate future advancements in the field.

## Introduction

In the past two decades, the study of microbiome composition and function has risen to the forefront of both medical and basic research (1, 2). In host-associated microbiomes, metagenomic whole-genome sequencing (WGS) has been widely deployed to show that microbiota composition and metabolic function have significant effects on the health of their host (1, 3, 4). The gut microbiome alone has been linked with a broad range of human diseases and disorders and is a target for therapeutic intervention (5, 6). Similarly, microbiomes found in water reservoirs (7), soil (8), and waste systems (9) were also found to play critical roles in modulating the chemistry of their respective environments. However, while such research primarily focused on prokaryotic microorganisms, viruses have also been shown to have major effects on microbiome dynamics (10, 11). Viruses are often the most abundant members of microbiomes and can play the roles of predators within an ecosystem through lytic activity, impacting population growth and nutrient turnover (12). Viral lysogenic activity can also affect community evolution through horizontal gene transfer events such as the spread of antimicrobial resistance (AMR) genes (13). These contributions of viruses to microbiome dynamics make them critical to study in both medical and basic research.

High-throughput metagenomic WGS of microbial communities allowed the study of viruses directly within their native environments, and advancements in viral sequence annotation led to a rapid expansion of viral sequence databases (14, 15). However, the assembly of complete viral genomes from metagenomes remained a major challenge due to their fast mutation rates and subsequent high heterogeneity in their sequences (16, 17). Long-read sequencing technologies from both Oxford Nanopore and Pacific Biosciences allow for the extraction of near-complete viral sequences (18, 19), however, such sequencing is prohibitively expensive compared to short-read sequencing of comparable depth and is typically only able to recover the most abundant viruses due to the lower number of reads (20). In prokaryotic genomes, this same challenge was overcome with the emergence of metagenomic binning software, which can extract genomes from short-read assemblies by predicting groups of contigs that belong to the same genomes. Coupled with methods to estimate the accuracy of such groupings with prokaryotic universal single-copy marker genes (21), such software has allowed the recovery of metagenome-assembled genomes (MAGs) from complex and diverse microbial communities (22–24).

Despite the widespread use and acceptance of metagenomic binning for prokaryotic genome recovery, similar advances have not been made for reconstructing viral metagenome-assembled genomes (vMAGs). One of the main challenges has been the inability to assess the completion and contamination of vMAGs from single-copy universal marker genes, as there is no such set known to exist for viruses(25). Several attempts have been made to bin viral contigs of select large phage genomes using conventional approaches (17, 26), however, the challenges of resolving closely related viral strains limit the broad application of this approach. Metagenomic binning approaches utilizing proximity-ligation sequencing (Hi-C, 3C, and other derivatives of chromosome conformation capture) show particular promise in reconstructing vMAGs (27, 28). Several studies have reported using the proximity-ligation signal to bin viral contigs together with their microbial host genome, with a high likelihood of these viral contigs belonging to the same viral genome (29, 30). Marbouty et. al remarked on using a custom application of the Louvain algorithm to reconstruct several possible vMAGs (31). However, these approaches rely on the assumption that each prokaryote may only host a single virus and that every virus may infect only one host (32), which is commonly not the case. To our knowledge, there is currently no available software designed for genome-resolved binning of vMAGs from fragmented metagenomic assemblies. The recent development of CheckV – software that can assess the completeness of viral sequences by comparing them to a large database of known viruses (33) enables such a software to be built and benchmarked.

Identifying the cellular hosts of viruses is critical for understanding their role in the microbiome, however, this information is lost during conventional shotgun or long-read sequencing, except for viruses integrated into the host genomes (prophages)(34). In the past, most virus-host association studies focused on probe- and emulsion-based interaction capture (35) and CRISPR array spacer alignment (36). However, proximity-ligated library sequencing allows for the most robust and high-throughput approach, as internalized viral DNA can be physically joined with the host chromosome DNA by *in vivo* crosslinking (36). One way of utilizing such data is to directly group viral contigs with their host genome during metagenomic binning with the proximity-ligation signal. This approach has been documented to produce robust virus-host associations (37, 38), but relies on the assumption that each virus can only infect one host, and is thus largely limited to prophages because more transient infections are likely to have weaker linkage signals (30). A second approach is to directly compare all viral sequences with all possible host MAGs to look for pairs with high proximity-ligation linkage signals, which allows for higher sensitivity and the identification of multiple host interactions from much weaker proximity ligation signal (39, 40). However, false-positive associations from poor library quality and read mis-alignments commonly found in proximity-ligation sequencing require robust linkage strength normalization and noise subtraction. To our knowledge, there is no software to perform such analysis, which has only been achieved on a sample-by-sample basis with custom methods. To overcome these gaps in the field and to accelerate high-throughput virus discovery and characterization, we developed ProxiPhage™ – a comprehensive end-to-end analysis platform for vMAG reconstruction and host prediction using proximity-ligation sequencing data.

## Results

### Viral MAG extraction improved genome completion

ProxiPhage is able to use proximity ligation sequencing to reconstruct viral genomes from highly fragmented short-read assemblies (Fig. 1). Viral contigs identified with VirSorter2 (41) in the assembly were grouped based on likely membership to original viral genomes with the viral binning function of ProxiPhage (Fig. 2; see Methods for details). In short, binning contigs based on Hi-C read linkages allowed for the extraction of groups of contigs that were likely in the same cell, phage envelope, or were otherwise in proximity with each other at the time of sampling. By comparison, binning the contigs using more conventional metagenomic metrics such as tetranucleotide frequency profiles and mean read coverages resulted in the grouping of viral contigs that are likely of similar phylogeny and abundance. Intersecting and resolving these two cluster sets allowed the reconstruction of viral metagenome-assembled genomes (vMAGs). The combined results avoid issues with the resolution of different viral genomes present in the same cell which would confound Hi-C deconvolution and similar codon usage profiles among viral families that would be poorly resolved by tetranucleotide frequencies. The synergistic combination of these two methods results in higher quality vMAGs than would be predicted by a simple merger of the two datasets.

**Fig. 1.**
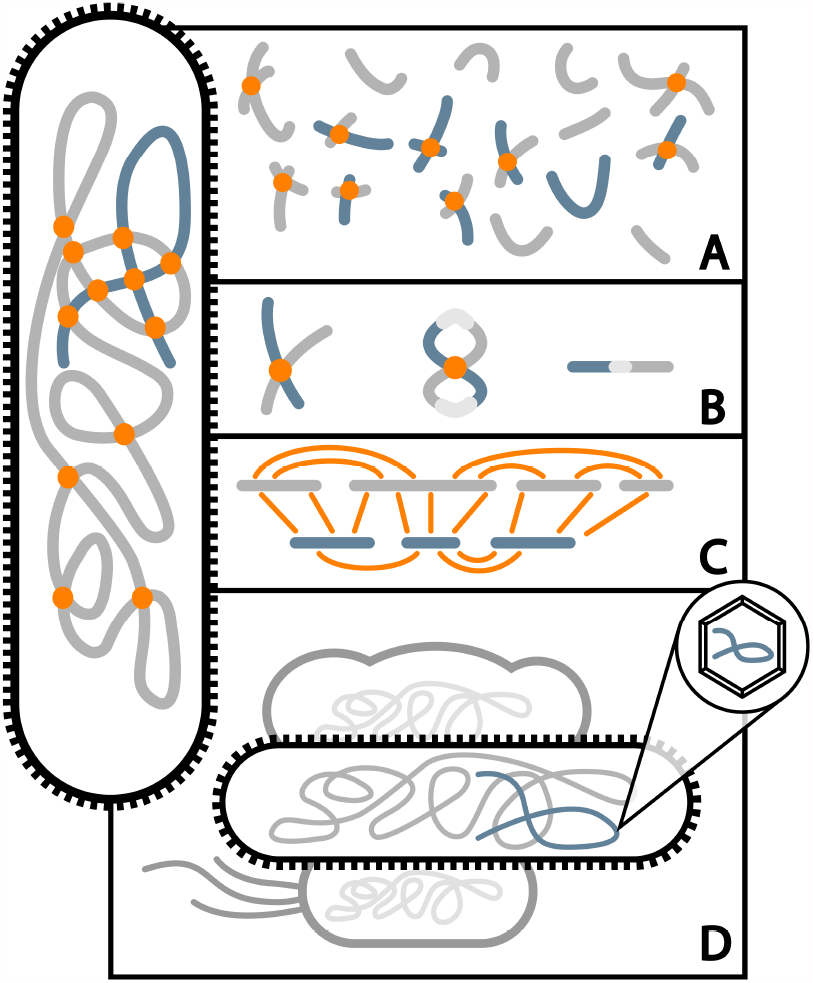
Proximity-ligation data use in ProxiPhage. Formaldehyde crosslinking *in vivo* physically constrains nearby DNA molecules inside the same cell (Left; host DNA in grey, phage DNA in blue). (A) Chromatin is fragmented to release crosslinked material containing DNA ends from nearby molecules. (B) Proximity ligation joins adjacent DNA molecules into chimeric junctions that are purified and sequenced. (C) The paired sequence information from chimeric junctions creates a connectivity matrix showing which contigs originated inside the same cells (including both phage-phage, phage-host, and host-host interactions). (D) Combined, this connectivity information can be used to associate phage and microbial contigs into MAGs and attribute viral MAGs to their microbial hosts within a mixed population.

**Fig. 2.**
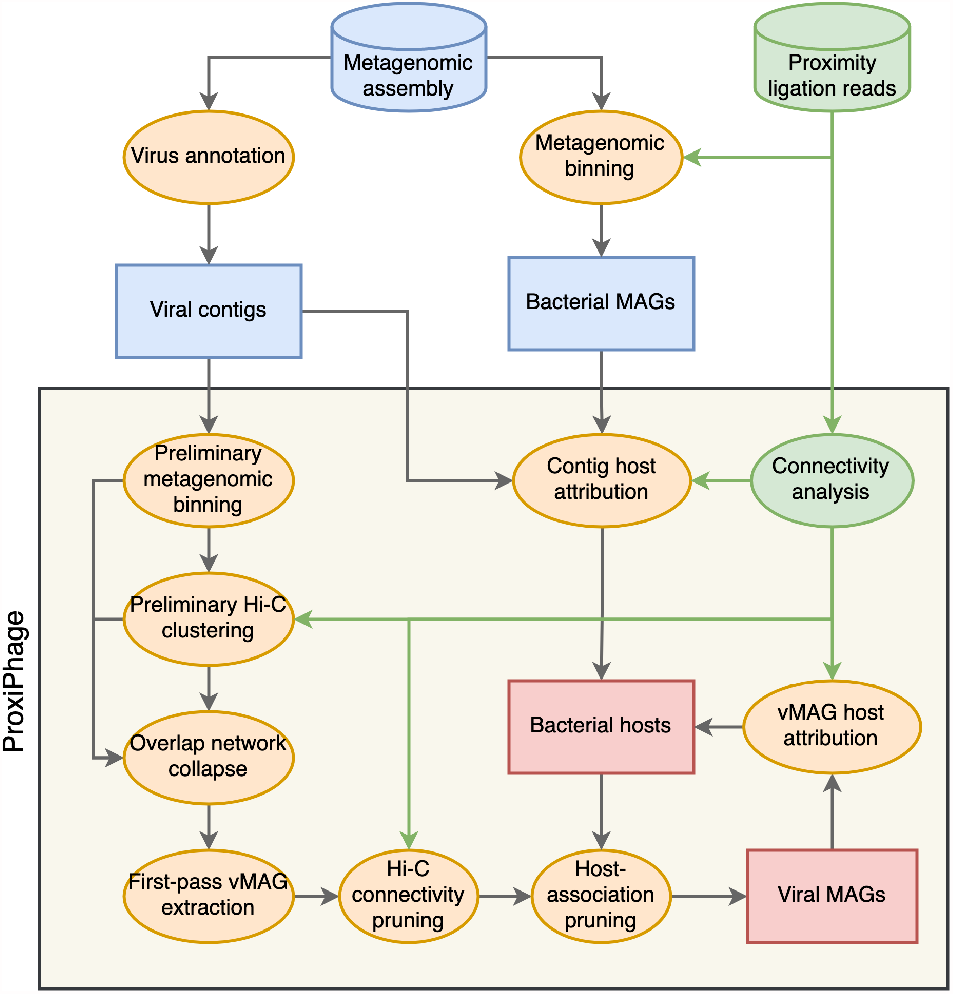
Computational pipeline. Flowchart showing the outline of data processing in ProxiPhage. Cylinders represent main input data, rectangles represent sequence data, and ellipses represent methods or software. Green represents Hi-C data and uses of the Hi-C data in the pipeline, orange represents the main components of the platform, blue represents input metagenomic data, and red represents the main ProxiPhage outputs.

To demonstrate ProxiPhage performance, we analyzed previously published proximity ligation sequencing data from a sheep fecal microbiome sample. In this metagenome, ProxiPhage was able to place 791 viral contigs into 315 vMAGs (Table 1). The sizes of the viral MAGs ranged from 11 - 197 kb in vMAGs consisting of 2-10 contigs each. vMAG binning increased the average length of predicted viral genomes from 18 kb to 45 kb and the final sequence N50 from 23 kb to 58 kb over that of the original set of viral contigs (Fig. S1). The genome completion of the ProxiPhage vMAGs was compared to that of the original viral contigs using CheckV (33), which uses an extensive viral lineage and protein database to estimate the completion of a given phage sequence. We found that the ProxiPhage vMAGs had significantly improved CheckV completion metrics (Fig. 3A). For instance, the number of near-complete viral genomes (>90% completion) improved from 9 to 73 after binning (Fig. 3C). For 276 out of the 351 (88%) of the vMAGs with a reliable reference, the completion improvements were also assessed using the alignments to the reference viral genomes from the long-read assembly (see below).

**Table 1.** Metagenomic data, assembly, binning and viral host attribution statistics from the four samples processed in this study.

| Category | Statistic | Sheep stool | Human stool | Cow rumen | Wastewater |
| --- | --- | --- | --- | --- | --- |
| General statistics | Hi-C reads (millions) | 100 | 100 | 100 | 95.3 |
|  | WGS reads (million) | 100 | 100 | 100 | 100 |
|  | Assembly size (Mb) | 873 | 479 | 531 | 822 |
|  | Assembly N50 | 4138 | 5922 | 3871 | 2511 |
|  | Number of contigs | 274,316 | 118,092 | 179,930 | 371,131 |
|  | Prokaryotic MAGs | 365 | 155 | 178 | 238 |
| Viral contigs | Number of viral contigs | 2341 | 1609 | 1298 | 1013 |
|  | Unfiltered virus-host links | 30,811 | 92,438 | 49,317 | 6561 |
|  | Filtered virus-host links | 1654 | 1314 | 1020 | 1024 |
|  | Viruses with hosts | 1526 | 1262 | 1000 | 705 |
|  | Percent viruses with hosts | 65.19% | 78.43% | 77.04% | 69.60% |
| Viral MAGs | Contigs in vMAGs | 791 | 714 | 599 | 340 |
|  | Number of vMAGs | 315 | 205 | 148 | 105 |
|  | Unfiltered vMAG-host links | 11,015 | 18,692 | 10,469 | 1538 |
|  | Filtered vMAG-host links | 244 | 185 | 125 | 126 |
|  | vMAGs with hosts | 216 | 180 | 122 | 87 |
|  | Percent vMAGs with hosts | 68.57% | 87.80% | 82.43% | 82.86% |

**Fig. 3.**
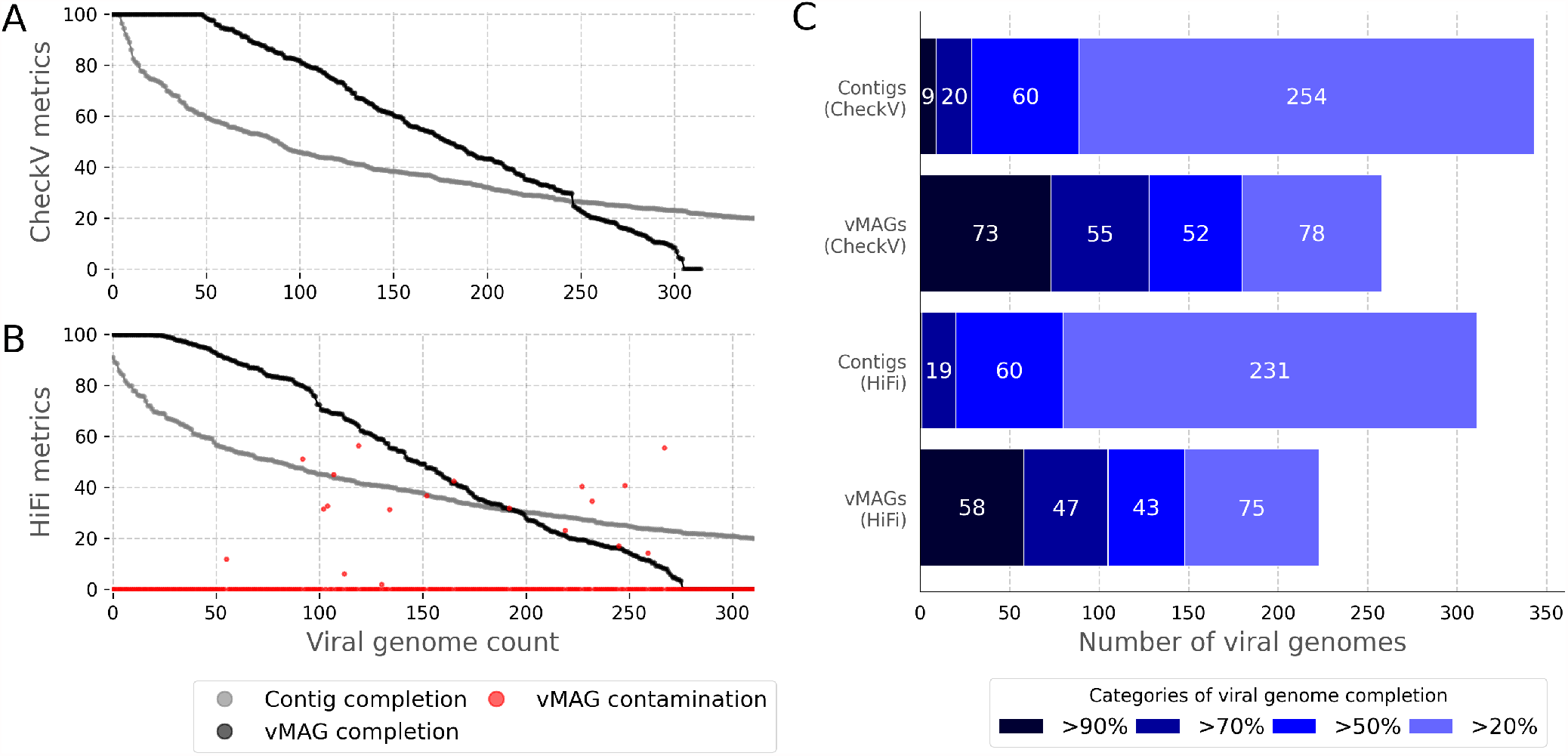
Viral MAG validation. The completion and contamination of unbinned viral contigs and binned vMAGs from a sheep fecal metagenome estimated with CheckV (A) and with viral references extracted from long-read HiFi assembly of the same sample (B), and a bar plot showing the number of high-completion viral genomes in the contigs and vMAGs (C).

### HiFi long-read assembly allows for vMAG validation

Due to the lack of a reliable universal marker gene set for viruses, the false-positive rates (or contamination) of the viral contig clusters must be evaluated with an orthogonal validation method. The sheep fecal microbiome sample described above was sequenced previously using PacBio HiFi chemistry (40). The resulting sequence data was used to generate a long-read assembly with much higher contiguity than the short-read assembly with an N50 of 279,621 bp compared to 4,138 bp, respectively. To serve as a reliable reference for both completion and contamination estimation, phage sequences were annotated and excised from the long-read contigs using VirSorter2 (see Methods). The resulting excised viral genomes from the HiFi assembly were assumed to be 100% complete and 0% contaminated for the purposes of validating vMAGs from the short-read assembly. The short-read and long-read phage sequences were aligned to each other and the similarity of the vMAGs and reference phages was assessed to estimate the percent completion and percent contamination of each vMAG that was present in the long-read assembly. To evaluate the validity of viral genome assessment using a long-read assembly, the estimated completion percentages of each phage contig and vMAG were compared to that estimated with CheckV. We found that both methods produced highly congruent completion scores (Figure S2).

### Viral MAGs are supported by the long-read assembly

Using the reference long-read virus sequences to estimate the completion of the vMAGs generated from the short-read assembly confirmed that the clustering method significantly improved viral genome completion (Fig. 3B) and increased the number of near-complete viral genomes from just 1 to 58 (Fig. 3C). The long-read references also allowed for the evaluation of vMAG contamination resulting from erroneous groupings of contigs that originated from different phages. We found that most of the contigs from any given vMAG aligned to a single long-read viral genome reference, confirming that the contigs originated from the same phage in the sample (Fig. 3B). However, there were still several instances of contigs from the same vMAG aligning to different reference phages. In the network visualization of select clusters (Fig. S3), possible contamination can be seen at vMAGs identifiers 23, 266, 140, 20, and 50. In total, we found that 17 of the 315 vMAGs (6%) had notable contamination of >10% (Table 2). The majority of these false positives were found to be closely related prophage sequences integrated in bacterial genomes of the fecal sample and thus could not be separated by the ProxiPhage algorithm.

**Table 2.** The number of vMAGs from a sheep fecal metagenome falling into broad quality categories after evaluation with reference viral sequences from a long-read HiFi assembly.

| Quality | Completion | Contamination | Count | Percent |
| --- | --- | --- | --- | --- |
| Near-complete | >90% | <10% | 57 | 20.65% |
| Moderate completion | >50% | <10% | 141 | 51.09% |
| Low completion | >0% | <10% | 259 | 93.84% |
| Over-contaminated | >0% | >10% | 17 | 6.16% |
| Testable | >0% | NA | 276 | 100.00% |
| Total | NA | NA | 315 | NA |

### Novel linkage signal normalization for evaluating virus-host interactions

ProxiPhage also features a novel approach for using proximity-ligation sequencing to infer the likely prokaryotic MAG hosts for viral contigs and vMAGs. The host finder independently evaluates each possible virus-host pair with at least 2 proximity-ligated read pairs linking them to estimate the average copy count of the virus genome per prokaryotic genome (see Methods; Formula 1). The density (links per *kb*^2^) of the virus-host is then normalized to the predicted virus per cell copy count and compared to the average intragenome Hi-C connectivity of the prokaryotic host (Formula 2) to evaluate the likelihood that the given MAG is the correct host for the virus. These normalization methods can be reliably used to threshold linkage data generated from a variety of Hi-C sequencing depths. The Hi-C sequences were down-sampled *in silico* to produce libraries ranging from 100 thousand to 100 million reads, and the resulting subsets were used to compute linkage metrics from the same virus-host pairs. Standardizing the linkage strengths by calculating the estimated viral copy count (Fig. 4A) and normalized connectivity ratio (Fig. 4B) revealed that these estimates do not significantly change with reduced Hi-C library depth. In addition to this, we also saw an enrichment of both these values around 1 – the theoretically expected value for both metrics in lysogenic infections (Fig. 4A-B; see below). Taken together, this suggests that these metrics can be used to reliably assess virus-host linkages regardless of the sequencing depth.

**Fig. 4.**
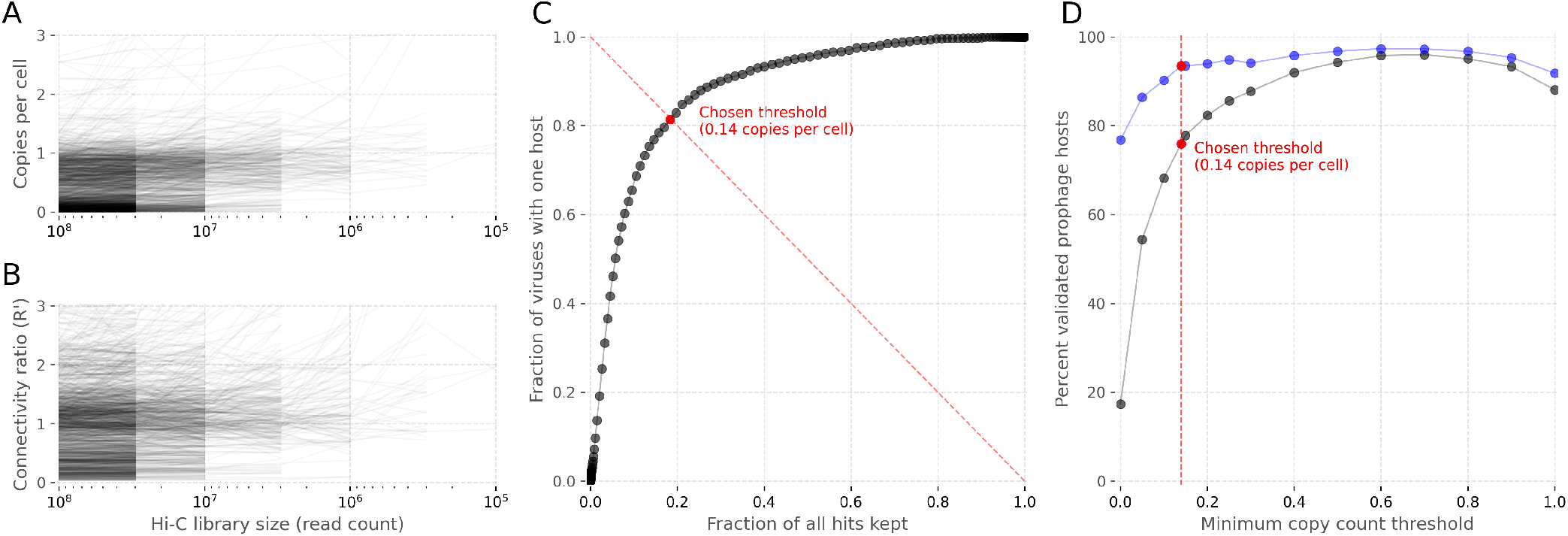
Host finding thresholding. Effect of Hi-C read rarefication (x-axis) on A) viral copy count per cell estimates and B) the normalized linkage ration R’. Each line represents one virus-host association tracked across multiple rarefied Hi-C libraries. C) Receiver operating characteristic curve showing the decline in the number of bacteria-phage host associations (x-axis) and in the number of phages with at least one host (y-axis) as the threshold of the minimum average copy count of each phage genome in its host is raised. D) The percent of prophage host links that were validated with HiFi long-read sequencing at different viral copy count thresholds. The red lines show the chosen threshold of 0.14 viral copies per cell.

### Unsupervised virus-host linkage thresholding

Evaluating the linkage strength based on the advanced normalization metrics allows for a more robust separation of true positives from false positives because the expected values are known. For the copy count metric, a value of 1 suggests that on average every cell in the population has 1 copy of the virus and thus is likely a true virus-host linkage. Likewise, a normalized linkage ratio of 1 means that the virus was connected to the host genome with the same signal strength as if it were part of the host genome.

The threshold chosen for the minimum copy count metric has a major impact on the number of interactions captured and the false-positive rate of the classifier. To set an optimal threshold, ProxiPhage automatically assesses the likely false-positive rates and false-negative rates at each cut-off and selects the optimal value based on the results (see Methods). In short, ProxiPhage constructs a receiver operating characteristic (ROC) curve and chooses a threshold that minimizes the fraction of the kept virus-host links while maximizing the number of viruses that still have at least one host (Fig. 4C). We observed that the area under the ROC curve (AUC) of this analysis and the chosen threshold can vary significantly depending on the quality of the proximity ligation library and the complexity of the sampled community. In the example of the highly complex sheep fecal microbiome analyzed in this study, the area under the curve (AUC) was relatively low – 0.88, and ProxiPhage selected a minimum copy count threshold value of 0.14 viral copies per cell.

To evaluate the accuracy of the automated thresholding and to validate this host-finding method, the long-read assembly was used to estimate the false-positive rates in the resulting associations. HiFi long-reads do not carry any inter-molecule information, so this validation was limited to prophages – viral sequences integrated into the host genome. The host sequence flanking prophages on the long-read contigs was compared to the sequence of the host MAG(s) that that the same prophage was linked to in the short-read assembly to determine if the host association was correct (see Methods). As expected, the true-positive virus-host assignment rate improved as the minimum copy count threshold was increased, peaking at 96% support (Fig. 4D, grey line). The automatically detected optimal threshold was placed at the point in the curve where further increasing the threshold started having diminishing returns and resulted in 731 prophage-host links with a true-positive rate of 74%. However, some of the false positives from this method could also be explained by the prophages being present in several hosts (but present only once in the long-read assembly). When the validation was rerun with prophages that were only assigned a single host (Fig. 4B, blue line), the automatically chosen threshold left 516 prophage-host links with a true-positive rate of 93%.

### ProxiPhage host attribution is sensitive and specific

The 2341 viral contigs identified by VirSorter2 in the sheep fecal metagenome sample were cross-referenced with 365 prokaryotic MAGs extracted from the same assembly using the ProxiPhage host attribution algorithm, resulting in a total of 30,811 possible virus-host pairs with at least 1 physical link (Fig. S4A). Applying multiple automated filtering steps to the data (see Methods) removed the vast majority of these links, leaving just 1654 links that were predicted to be true positives (Fig. S4B, Fig. 7A). In total, the algorithm was able to identify a prokaryotic host for 1526 (65%) of the viral contigs. Of these, 588 were found to be prophages with a single host (see Methods), with a true host attribution rate of 93% as validated with the long-read reference assembly. While the majority of the viral contigs were assigned just one host, several viral contigs were linked to multiple prokaryotic MAGs, suggesting possible promiscuous viruses (Fig. S4, vertical lines). Likewise, some prokaryotic MAGs were linked with many viral contigs, which could be assembly fragments of the same viruses or indicate co-infection (Fig. S4, horizontal lines). Reassuringly, the majority of contigs that were clustered together into a vMAG were assigned identical or similar prokaryotic hosts (Fig. 5), except where coverage dropout caused likely false negatives. It should be noted, however, that some of the weaker host associations still appear to be clear outliers in their respective vMAG clusters, indicating the presence of residual false-positive associations even after thresholding.

**Fig. 5.**
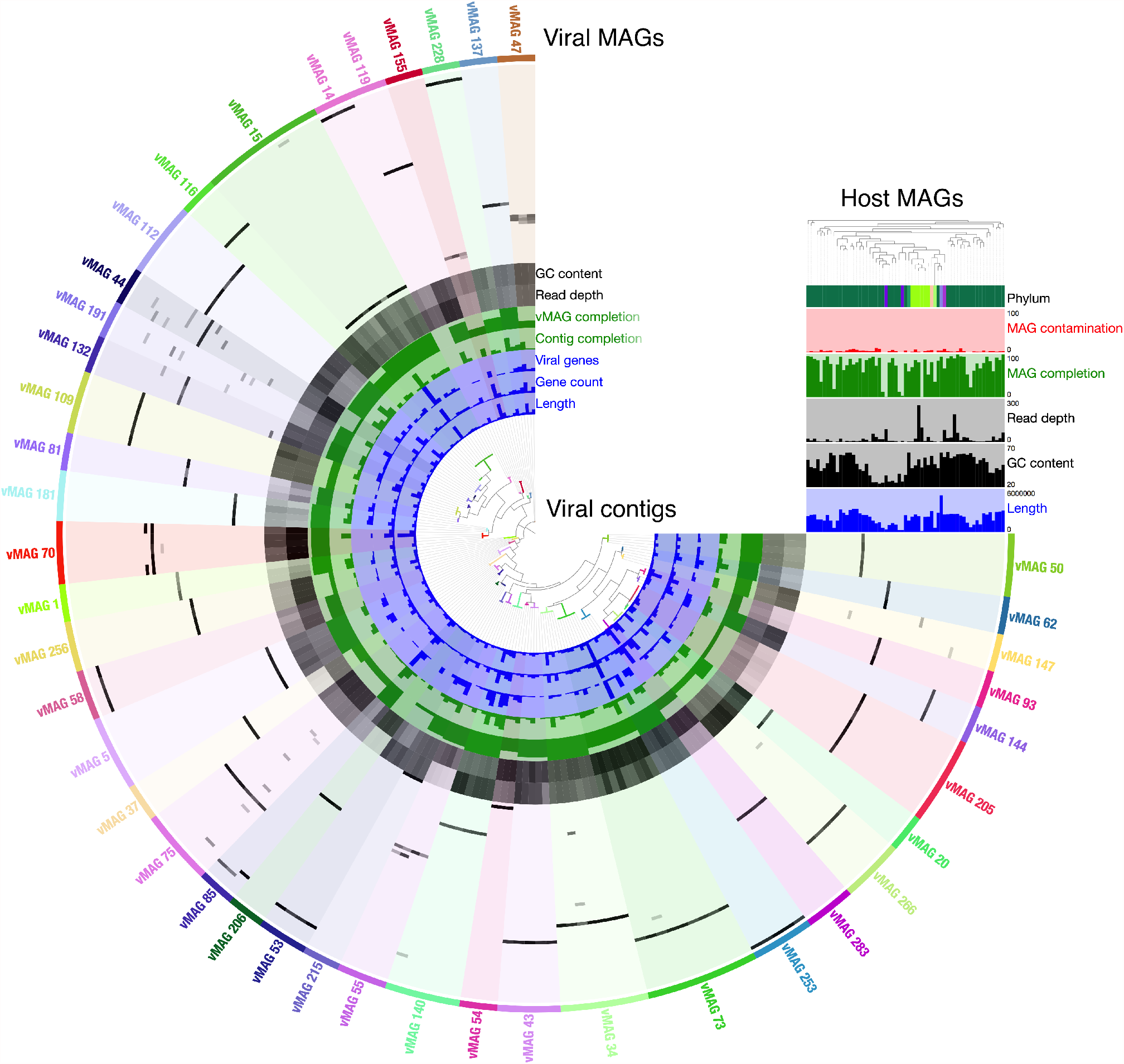
Viral MAGs and their hosts. Viral metagenome-assembled genomes (vMAGs, outer ring) derived from a sheep fecal metagenome, with their associated contigs in the same color. Darker bars indicate higher estimated copy count. Additional circular layers (viral) and bar plots (prokaryotic host MAGs, upper-right) represent characteristics of viral contigs or host MAGs respectively, including gene content, length, GC%, and estimates of completion based on alignment to a long-read metagenomic assembly. Radial grey bars indicate a viral-host association, with the intensity encoding estimated viral copy count per cell. Only vMAGs with at least 3 contigs and a host found for every contig are shown to fit this visualization.

**Fig. 6.**
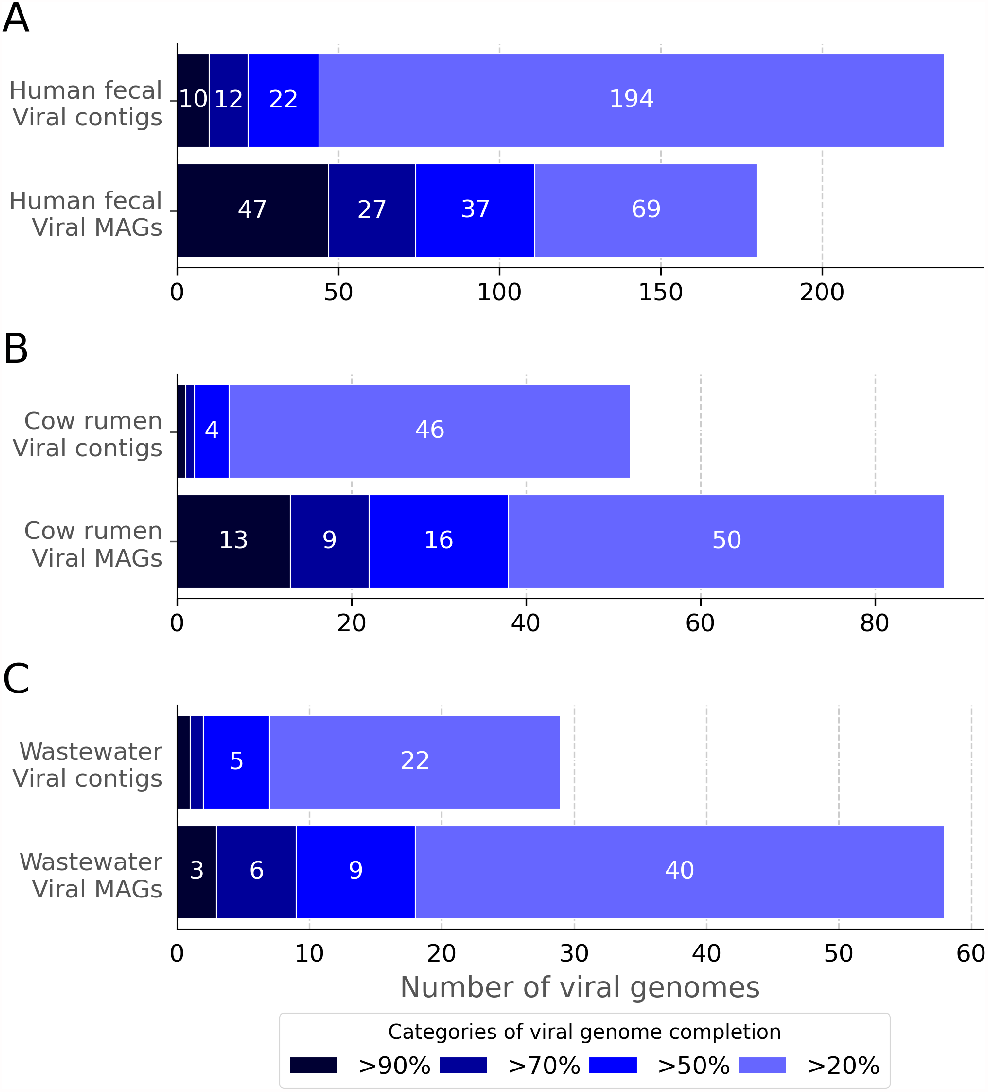
Viral MAG extraction in additional samples. Viral genome completion (estimated with CheckV) of the original and binned viral contigs from additional benchmarking metagenomic samples extracted from A) human stool, B) cow rumen, and C) wastewater. Bar plots show the number of genomes at different completion cut-offs in the original viral contigs and vMAGs, as estimated with CheckV.

**Fig. 7.**
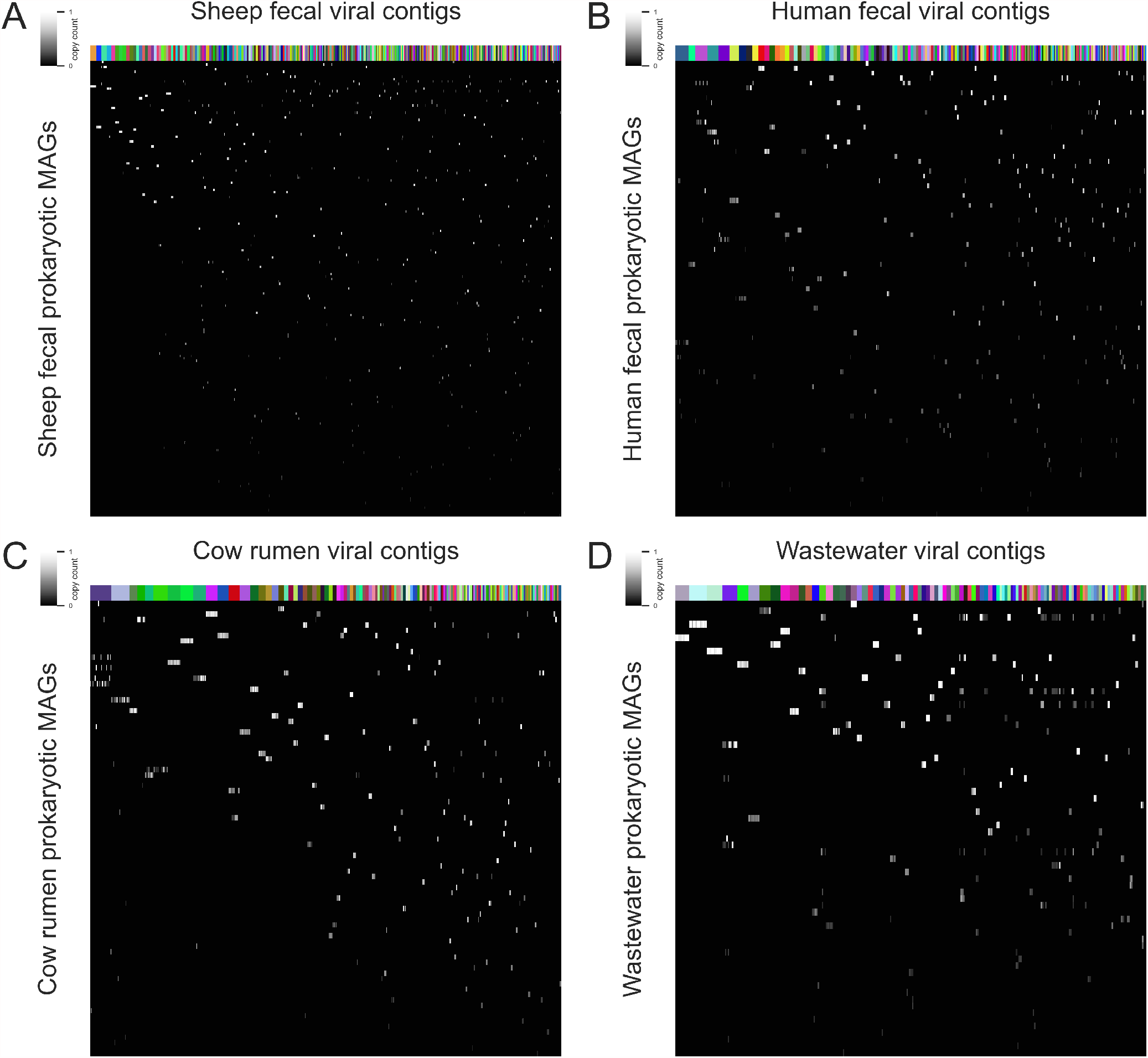
Viral host assignment in additional samples. Prokaryotic hosts identified for viral contigs with ProxiPhage from additional benchmarking microbiome samples extracted from A) sheep fecal, B) human stool, C) cow rumen, and D) wastewater. The color map encodes for the estimated average copy count of each phage genome in its host. Columns are clustered according to vMAG membership (labeled with random colors) and rows are grouped based on linkage similarity with seaborn clustermap. Only viral contigs from viral MAGs are shown.

The host attribution algorithm in ProxiPhage works more reliably on complete vMAGs since the algorithm has more sequence length to use for estimating the connectivity likelihood. When using the vMAGs for host assignment, applying the automatic filtering on the unfiltered connectivity matrix (Fig. S5A) results in a cleaner matrix, with relatively few predicted promiscuous phages or host co-infections (Fig. S5B). Interestingly, the algorithm predicted the optimal copy count value to be 0.14 – the same as in the original viral contigs. This reduced the possible 11,015 vMAG-host links with at least 1 Hi-C link to just 244 links after thresholding, which still associated a total of 216 vMAGs (69%) with a prokaryotic MAG host.

### ProxiPhage can resolve viral genomes and their hosts from a variety of sample types

The sheep fecal microbiome used for benchmarking the accuracy of de-novo viral binning and host finding is the only currently available sample that has been sequenced to such a high depth with long-read sequencing (40). To test the applicability of ProxiPhage to a variety of medically and environmentally relevant microbiome sample types, we processed a fecal sample from a healthy human donor, a cow rumen sample, and a sample from a wastewater treatment plant (see Methods). Using the same computational analysis pipeline as the sheep fecal sample, we assembled and annotated a total of 1609, 1298, and 1013 viral contigs from these samples, respectively (Table 1, S1). These viral contigs were then binned into 205, 148, and 104 vMAGs in each sample, and their completion was compared to that of the original viral contigs with CheckV (Fig. 6). We found that in every sample, ProxiPhage significantly increased the completeness of the resulting viral genomes. The impact of viral binning was particularly notable in the human fecal and cow rumen samples, where many of the vMAGs had 10 - 27 contigs. The metagenomic assemblies from the human fecal, cow rumen, and wastewater samples were also binned into prokaryotic MAGs to test the host attribution pipeline. The viruses from these three samples were linked to the MAGs of their respective communities using the proximity ligation signal to identify their likely cellular host genomes. The application of automated minimum copy count filtering on the three samples resulted in significantly different thresholds being chosen – 0.03, 0.03, and 0.12 copies per cell, respectively. For the human fecal and cow rumen sample, the AUC of the ROC analysis was 0.97 and 0.98 viral copies per cell, respectively, suggesting a very high signal-to-noise ratio in the Hi-C linkage data, while the AUC for the wastewater sample was lower, at 0.86. After final thresholding, ProxiPhage retained 1314, 1020, and 1024 high-quality virus-host links, and a total of 1262 (78%), 1000 (77%), and 705 (69%) viruses were assigned at least one host in the human fecal, cow rumen, and wastewater samples, respectively (Fig. 7; Table 1). Similar to the results from the sheep fecal sample, most of the viral contigs from the same vMAGs were assigned to the same host(s), confirming the accuracy of viral bin and host assignments (Fig. 7, top color bar).

## Discussion

ProxiPhage is the first automated software capable of accurate extraction of near-complete viral genomes from highly fragmented metagenomic assemblies by using proximity-ligation sequencing such as Hi-C. This approach is significantly cheaper and more scalable for large-scale studies compared to long-read sequencing of comparable depth, while also enabling the association of the viruses with their hosts (20). Viral binning has been previously indirectly achieved with custom analysis of select samples by placing multiple viral contigs together with their host genome sequences (29–31). This approach can fail in events of promiscuous phage infections (32, 42, 43), which have been observed in all four microbiome samples investigated in this study. By sorting viral contigs with both proximity-ligation signal as well as conventional binning methods, ProxiPhage allows for viral genome-resolved de-convolution even in events of phage co-infection, promiscuous phages, and relatively low Hi-C coverage. Since the majority of virus discovery efforts focus on extracting viral genomes from short-read metagenomic assemblies (44) which often results in recovering short, fragmentary contigs (17, 45), ProxiPhage has the potential to greatly accelerate the discovery of near-complete viral genomes.

Proximity ligation sequencing has the added benefit of capturing interactions of viruses with their respective hosts. In both environmental and medical applications, capturing this information is crucial for understanding the impact and contribution of viruses on the functioning of their respective communities (46). The high throughput and relatively low cost of proximity ligation sequencing make it the most scalable approach for use in untargeted virus association and characterization studies (36, 39). Several studies have been able to predict the prokaryotic MAG hosts of viruses using this data type, however, these studies relied on careful investigation and custom thresholding to produce their results. ProxiPhage offers an unsupervised approach for such analysis and overcomes many challenges of proximity linkage data normalization and thresholding, making it appropriate for use on a variety of sequencing depths, library qualities, and community compositions. As demonstrated in the four metagenomes processed in this study, ProxiPhage yields robust virus-host associations for the majority of viruses, making it appropriate for large-scale viral infection screening studies. In addition to a binary host association output, the provided normalized metrics can provide useful information about the nature of a virus-host interaction. The average copy count of the virus per host cell can help distinguish relatively rare infection events, common sample-wide associations, or even active phage replication in the host cells. On the other hand, the normalized linkage ratio indicates how closely associated the viral DNA is with the host chromosome(s), allowing the identification of integrated prophage sequences.

Neither the binning nor the host-attribution features in ProxiPhage rely on a priori sequence modeling, suggesting that it may also be applied for the study of other mobile genomic elements, such as plasmid sequences or gene cassettes. However, this functionality was not shown or benchmarked in this study, primarily because the sheep fecal metagenome used for benchmarking was found to have very few plasmid sequences. The challenges of plasmid genome binning and host association are, in principle, very similar to those of prophage analysis (38), and plasmid binning and host attribution are very difficult to achieve without proximity-ligation data (47, 48). Similar to viruses, plasmid sequence reconstruction (49) and host association (27, 28, 38) have been previously achieved using proximity-ligation sequencing, however, ProxiPhage is the first unsupervised software capable of this function. The possibility of high throughput plasmidome de-convolution could have a great impact on microbial resistome characterization in a variety of research and medical fields (48).

The automated vMAG extraction and host attribution features of ProxiPhage make it the first software capable of such analysis, and its accuracy and high throughput have the potential to aid the study of viruses and their cellular hosts. In both environmental and host-associated metagenomic studies, this platform can accelerate the discovery of novel viral clades and improve our understanding of the role of phages in the composition dynamics and nutrient cycling of their respective communities (50). Finally, our platform could be applied in clinical settings for efficacy and safety screenings of fecal microbiota transplantations (FMT) and phage therapies, and for predicting the effects of such treatments on specific patients (51, 52).

## Methods

### Read and assembly pre-processing

DNA extracted from sheep feces was processed and sequenced as described in Bickhart et. al (40). To better represent common sequencing depth and for consistency with other samples in this study, the shotgun sequences were downsampled to 100 million reads and then assembled with MegaHit (53) v1.2.9 using default parameters. The resulting assembly had 274,316 contigs at least 1 kb in length and contained a total sequence length of 873.4 Mb with an N50 of 4138 bp (Table 1). A proximity ligation library was also prepared from the same sample using the ProxiMeta™ Hi-C kit from Phase Genomics, and the resulting 2×150 bp sequences were also downsampled to 100 million reads. These Hi-C reads were then aligned to the metagenomic assembly with BWA (54) v0.7.17 and compressed with Samtools (55) v1.10. The alignment was then scanned to count the total number of long-range physical (Hi-C) interactions between contigs (different contigs or on the same contig but at least 10 kb apart). Similarly, the number of close-range interactions was also counted and saved (at most 10 kb apart on the same contig). Only alignments that were non-redundant, full-length, and with a maximum of 1 mismatch were considered in these tallies. A total of 14,597,959 long-range and 44,233,451 close range Hi-C interactions were recorded. Finally, a size-selected SMRTbell (56) library (9-14 kb final fragment length) was prepared from the same sample for ultra-deep sequencing on the Sequel, yielding a total of 255 Gb of long-read HiFi data, as described in Bickhart et. al (40). These long reads were then assembled with metaFlye (57) to produce 60,050 contigs with a total of 3.43 Gb of sequence and an N50 of 279,621 bp. This HiFi assembly was used as a reference to evaluate binning and host association methods in this paper.

### Viral sequence annotation

Long contigs (>5 kb) in both the short-read and long-read assemblies were annotated with VirSorter (41) v2.2.2 with default parameters to find likely viral sequences. For the shortread assembly, original unmodified contigs with at least 50% viral gene content were saved for subsequent downstream analysis. For the Hi-Fi long-read assembly, predicted viral genomes were excised with VirSorter2 and used as complete viral genome references. Using these methods, 2341 (N50 15,919 bp) and 8054 (N50 66,873 bp) viral sequences were annotated in the short-read and long-read assemblies, respectively.

### ProxiPhage viral binning

ProxiPhage viral binning features a combination of proximity ligation signal clustering and conventional metagenomic binning approaches to overcome the limitations of either method. First, the viral sequences are binned with both methods to produce two preliminary sets of vMAG. A non-redundant set overlap network is constructed from these two contig groupings such that nodes represent contig clusters from either of the two preliminary vMAG sets, and edges contain contigs that overlap between bin sets. These overlaps are then scanned and resolved through a proprietary greedy network collapse algorithm featured throughout the ProxiMeta (38) platform to produce a single set of vMAGs that is more accurate and complete than either of the original inputs. Each vMAG cluster is then additionally scanned and adjusted to ensure a minimum strength of Hi-C linkages between its contained contigs, which further reduces possible contamination. Finally, ProxiPhage viral binning also allows for additional pruning of vMAG clusters such that contigs from each cluster have similar or identical predicted prokaryotic hosts, although this feature was not utilized in this study since host commonality was one of the metrics used for vMAG accuracy assessment.

### Viral MAG validation and benchmarking

The viral genome completion of both viral contigs and vMAGs was estimated with CheckV (33) v0.7.0 with default parameters. Note that CheckV does not natively support the investigation of contig clusters, so to run CheckV on the vMAGs the sequences from each vMAG needed to be concatenated into a single sequence with 200 bp “N” spacers. Also note that the “contamination” metric produced by CheckV refers to the bacterial content of the sequences, and not binning false positives as it does CheckM (21). The quality of the viral contigs and vMAGs was also assessed by comparing the sequences to the reference phage sequences excised from the long-read HiFi assembly (40). The short-read and long-read phage sequences were aligned to each other using BLAST (58) v2.11.0, and high-quality alignments (>95% percent identity, >100 bp length) were saved. The reference alignment network was constructed from these alignments using Cytoscape (59) v3.7.1. The best reference for each vMAG was determined as the reference to which the greatest percentage of its sequence aligned, with a minimum of 1 kb. The completion of the vMAG was estimated as the percentage of the reference that aligned to the vMAG (Formula 1), and its contamination was estimated as the percentage of the vMAG sequence that did not align to the reference (Formula 2). Contigs or contig segments that did not reliably align to any of the references were not counted in the contamination calculation to account for some of the short-read assembly sequences not being present in the long-read assembly.

Genome completion of a query viral sequence *ω*(*q*) calculated using a long-read reference assembly from the total length of the reference that aligned to the query *A*(*r*) and the length of best reference genome *L*(*r*):

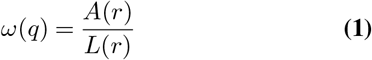

Genome contamination *χ*(*q*) of a query viral sequence calculated using a long-read reference assembly from the total length of the query *L*(*q*, the length of the query that aligned to the best reference *A*(*q*) and the length of the query that was not found to align to any reference sequence *L*(*u*):

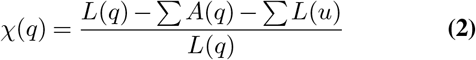

### Prokaryotic MAG extraction

The metagenomic assembly was de-convoluted with the ProxiMeta (38) platform to extract draft prokaryotic genomes to be used for finding the likely hosts of the viral sequences. The completion and contamination of the MAGs were estimated with CheckM (21) v1.1.3. In total, 365 MAGs were formed, representing 34% of the total assembly sequence. Of these, 151 MAGs were of moderate quality (>50% completion and <10% contamination), and none of the 365 MAGs were over-contaminated (contamination >10%). All the resulting clusters were used for viral host attribution with ProxiPhage.

### ProxiPhage host attribution

The long-range Hi-C linkage data was scanned to identify viral contigs and prokaryotic MAGs with a Hi-C link between them. A combination of the Hi-C link count, viral read depth, and MAG read depth were then used to estimate the average copy count of each virus in each MAG (Formula 3). The density of Hi-C links per *kb*^2^ of sequence between the virus and the MAG was then compared to the connectivity of the MAG to itself and normalized to the estimated copy count to compute the normalized connectivity ratio (Formula 4). This value assesses the strength of the virus-host linkage in the context of what would be expected if the virus was found inside the cell, with a value of 1 being ideal. Virus-host linkages were then filtered to keep only connections with at least 2 Hi-C read links between the virus and host MAG, a connectivity ratio of 0.1, and intra-MAG connectivity of 10 links to remove false positives. For the final threshold value, a receiver operating characteristic (ROC) curve is used to determine the optimal copy count cut-off value. The optimal cut-off was determined from the ROC curve as the value that produces the point to the top left of the plot, or the cut-off that removed the maximum number of virus-host links while still finding at least one host for the maximum number of viruses. Each virus is also evaluated for the fraction of host MAGs that it still had connections with to identify “sticky” sequences with a likely high proportion of false positives. These were corrected by removing linkages with an average copy count less than 80% of the highest copy count value for the given viral sequence. The above host assignment workflow also works identically for vMAG clusters, but with linkage and length values from different contigs being added together.

Average viral copy counts per cell *C* calculated from the virus abundance *V*, prokaryotic host abundance *H*, Hi-C links between the virus and host *L*, and total Hi-C links of the virus and all possible hosts *L*(*v*):

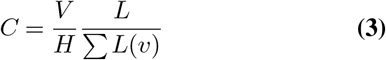

Normalized connectivity ratio *R*′ calculated from the Hi-C connectivity density between the virus and host *D*_*V H*_ and of the host genome to itself *D*_*H*_, and normalized to the virus abundance *V*, prokaryotic host abundance *H*, Hi-C links between the virus and host *L*, and total Hi-C links of the virus and all possible hosts *L*(*v*):

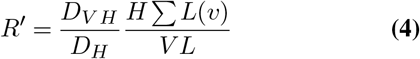

### Prophage host validation

The accuracy of prophage virus-host links found with the host attribution software in ProxiPhage was evaluated with the reference long-read HiFi assembly. The short-read viral sequences were aligned to the full HiFi assembly using Blast (58) v2.11.0, and high-quality alignments (> 95% percent identity, > 100 bp length) were saved. The best reference contig for each virus was defined as the contig to which the greatest percentage its sequence aligned to, with a minimum of 50%. If the HiFi contig still had more than 200 kb of sequence that did not align to the viruses, the virus was declared a prophage and thus used for subsequent analysis. This unaligned bacterial sequence was compared to the sequence of the host MAG from the short-read assembly to which the virus was linked. If at least 10 kb of the host sequence aligned to the host MAG, the virus-host link was considered correct. Only good quality MAGs (>50% completion, <10% contamination according to CheckM) were included in this analysis.

### Anvi’o analysis

A custom selection was taken from the full vMAG set for detailed visualization with Anvi’o 7 (60). This selection contained vMAGs that had at least 3 contigs and had at least one host assigned to each of its contigs. The phylogenetic tree was constructed from the vMAG sequences using VICTOR (61), and the resulting Newick tree was then manually modified to replace the vMAG leaves with the contigs contained in each of the vMAGs. For the host MAGs, their taxonomies at the phylum level were estimated manually using a combination Kraken2 (62) v2.1.1 (using default options and the full standard database) and metaWRAP (63) v1.3.2 blobology and *classify*_*b*_*ins* modules using default options. The phylogenetic tree of the host MAGs was then constructed with Anvi’o 7 (60) using a concatenated alignment of all common ribosomal genes. The estimated average read depth for both viral contigs and host MAGs was estimated with the *jgi*_*s*_*ummarize*_*b*_*am*_*c*_*ontig*_*d*_*epths* script in MetaBAT (22) v2.15.

### Human fecal microbiome processing

Libraries were prepared using the Phase Genomics ProxiMeta Hi-C kit version 3 following the manufacturer’s protocol from a healthy donor stool sample. In brief, approximately 250 mg of sample was homogenized in phospho-buffered saline (PBS) and pelleted by spinning at 17,000 x g for 1 min. The pellet was resuspended in 1 ml of crosslink solution and incubated at room temperature for 20 min at room temperature with rotational mixing. Crosslinking was terminated by the addition of 100 µl of Quench solution and incubation for 15 min at room temperature with mixing. After pelleting sample at 17,000 x g for 5 minutes, pellets were washed once in chromatin rinse buffer (CRB) and then resuspended in 700µl of Phase Genomics Lysis buffer 1 and 250 µl of Lysis beads. The sample was placed in a Turbomix disruptor (Scientific Industries) and mixed at maximum speed for 20 minutes. The lysate was spun down briefly, and the lysate was transferred to a new microcentrifuge tube and chromatin pelleted by spinning at 17,000 x g for 5 minutes. The pellet was then washed with CRB and resuspended in 100 µl of Phase Genomics Lysis Buffer 2 and incubated at 65°C for 15 min. Chromatin was then bound to Recovery beads, washed with CRB, and then fragmented/ends filled in with biotinylated nucleotides at 37°C for 1 h. Beads were washed and resuspended in 100 µl of Proximity Ligation buffer and 5 µl of Proximity Ligation Enzyme and incubated at 25°C for 4h. Reverse crosslinks enzyme was added at the sample was heated to 65°C for 1h to release DNA from crosslinked chromatin. DNA was purified using Recovery Beads and biotinylated ligation junctions capture using streptavidin beads. Bead-bound DNA was used to generate a dual unique-indexed Illumina-compatible library. DNA for Shotgun WGS libraries were prepared using a ZymoBiomics DNA miniprep kit (Zymo Research). Shotgun libraries were prepared using Nextera XT (Illumina) following the manufacturer’s protocol and 50 ng of input DNA.

## Additional sample processing

Additional benchmarking data was generated from the short-read WGS and Hi-C sequencing data from the cow rumen metagenome (64), the wastewater benchmark metagenome (48), and the human fecal metagenome (see human fecal microbiome processing). All downstream sequence analyses, including rarefaction to 100 million reads, metagenomic assembly, viral annotation, viral binning, and host attribution were identical to the analysis performed on the main sheep metagenome (Table 1).

## Data Availability

The raw data from the sheep fecal shotgun WGS and Hi-C sequencing is publicly available from NCBI BioProject PRJNA595610. The cow rumen microbiome data is available from BioProject PRJEB21624, sample SAMEA104567052. The wastewater microbiome data is available from BioProject PRJNA506462. The human fecal microbiome data is available at https://proximeta.phasegenomics.com/proximeta-pgfecal. The four shotgun assemblies used in this study are available at https://bitbucket.org/phasegenomics/proxiphage_paper/src/main/assemblies. The sheep fecal microbiome long-read HiFi assembly used as a reference in this study is available at DOI: https://doi.org/10.5281/zenodo.4729049. All other data, intermediate files, and analysis scripts are publicly available from https://bitbucket.org/phasegenomics/proxiphage_paper. The ProxiPhage viral analysis platform is available through the ProxiMeta™ service platform at https://proximeta.phasegenomics.com.

## Acknowledgements

We thank the early users of ProxiPhage for their patience and feedback during testing and for their suggestions for the platform’s features. We also thank the engineering and management teams of Phase Genomics for their support, help, and code review during ProxiPhage development and the laboratory staff for constructing sequencing libraries. We also wish to thank Natalie Solonenko and Marie Burris for helpful technical discussions.

## Author Contributions

GU conceived, developed, and benchmarked ProxiPhage, processed the results, and wrote the manuscript; MP developed the prototype of the viral host attribution algorithm; CS, GDH, and AZ assisted with developing experiments and data analysis, AW and JG deployed and maintained the ProxiPhage cloud service; DB and TS provided benchmarking sequence data, assemblies, and taxonomic annotations; SS and IL directed ProxiPhage development; BA, SE, and MS provided guidance and insight for the project; IL and GU conceived the project. All authors read and edited the manuscript.

---

**Fig. S1.**
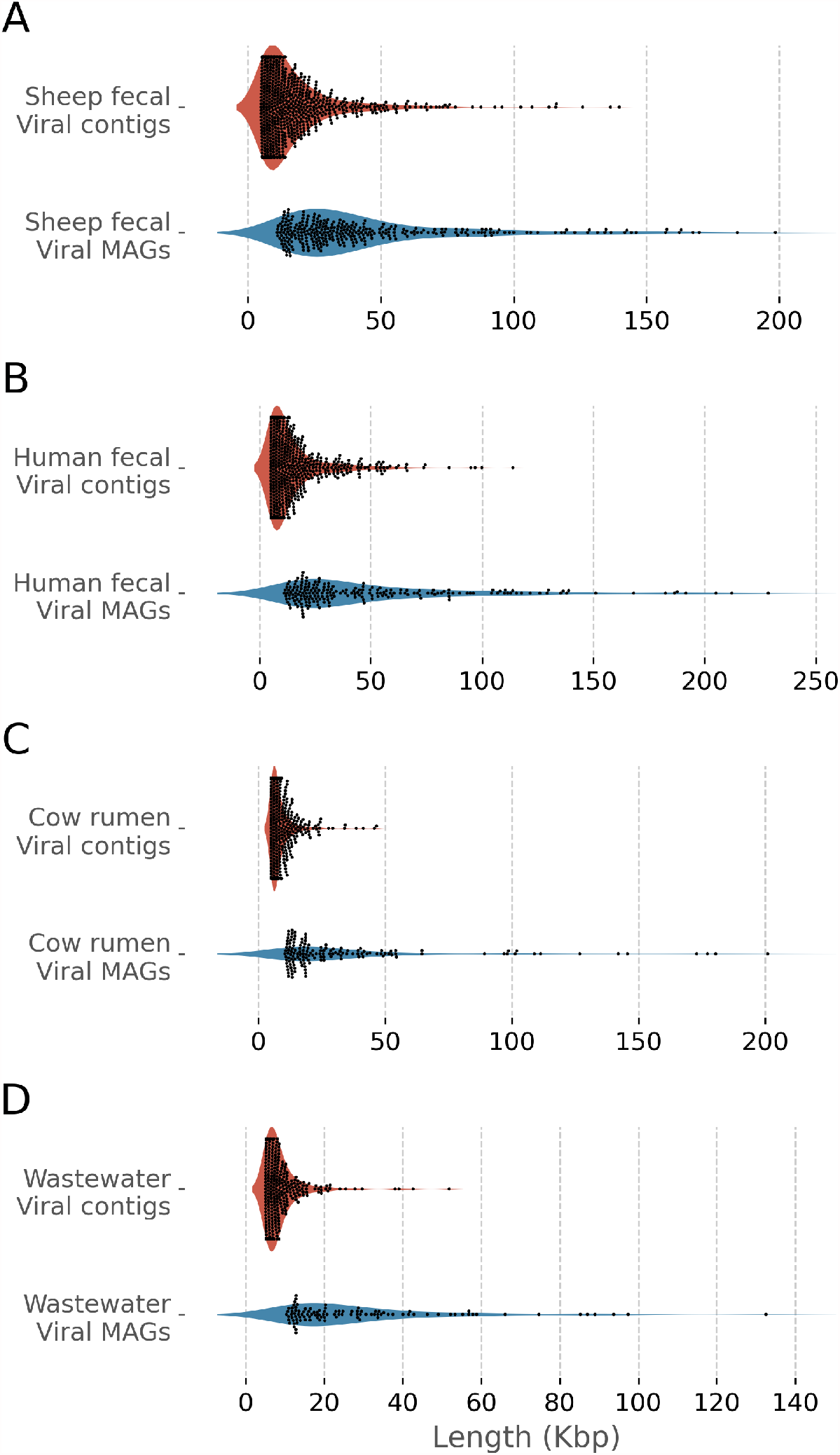
Viral MAG lengths. The length distribution of viral sequences before (red) and after (blue) binning with ProxiPhage in metagenomic samples extracted from A) sheep stool, B) human stool, C) cow rumen, and D) wastewater.

**Fig. S2.**
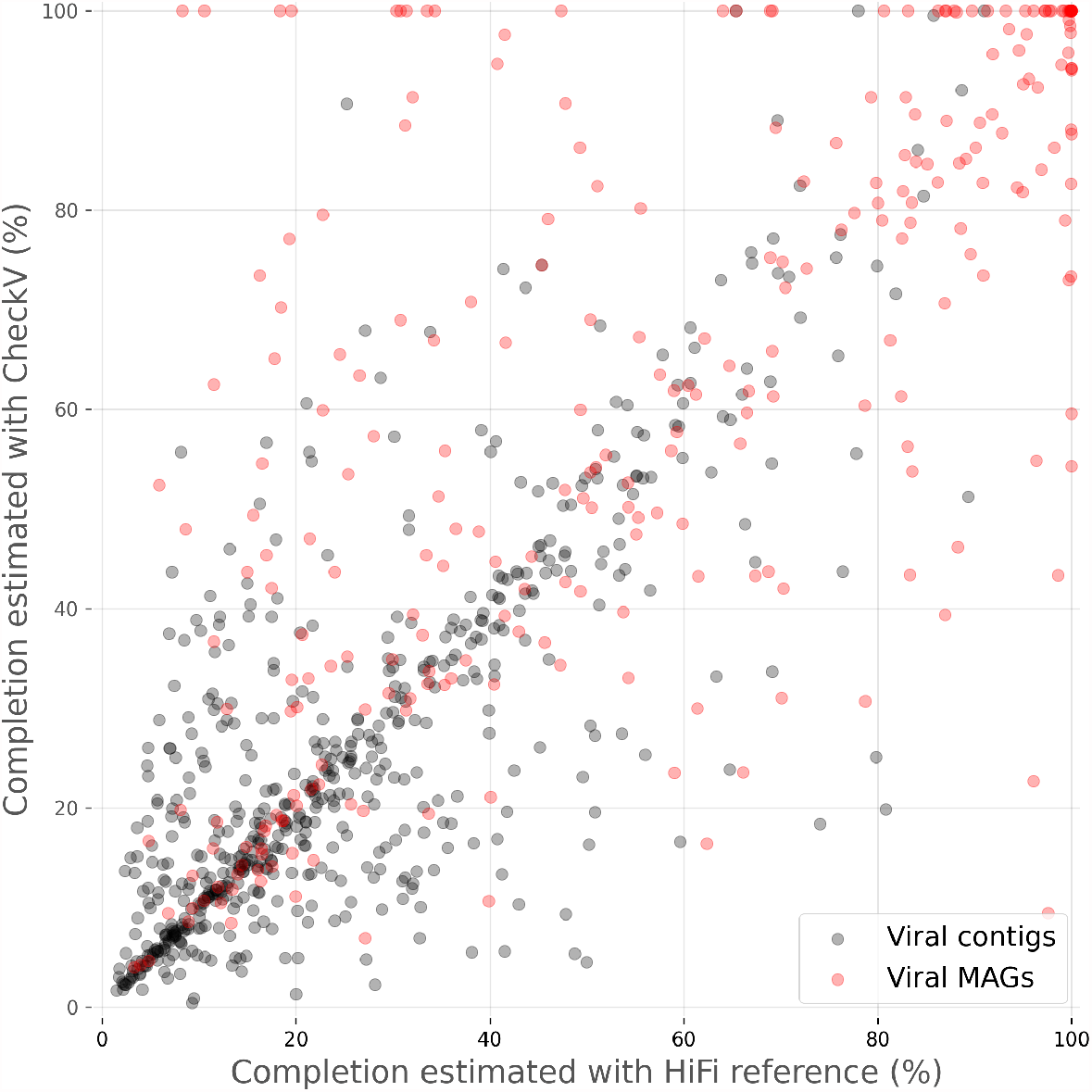
Completion estimation comparison. Scatter plot of completion percentages of viral contigs and vMAGs from a sheep fecal metagenome, estimated with CheckV (y-axis) and with reference excised phages from a long-read HiFi assembly (x-axis).

**Fig. S3.**
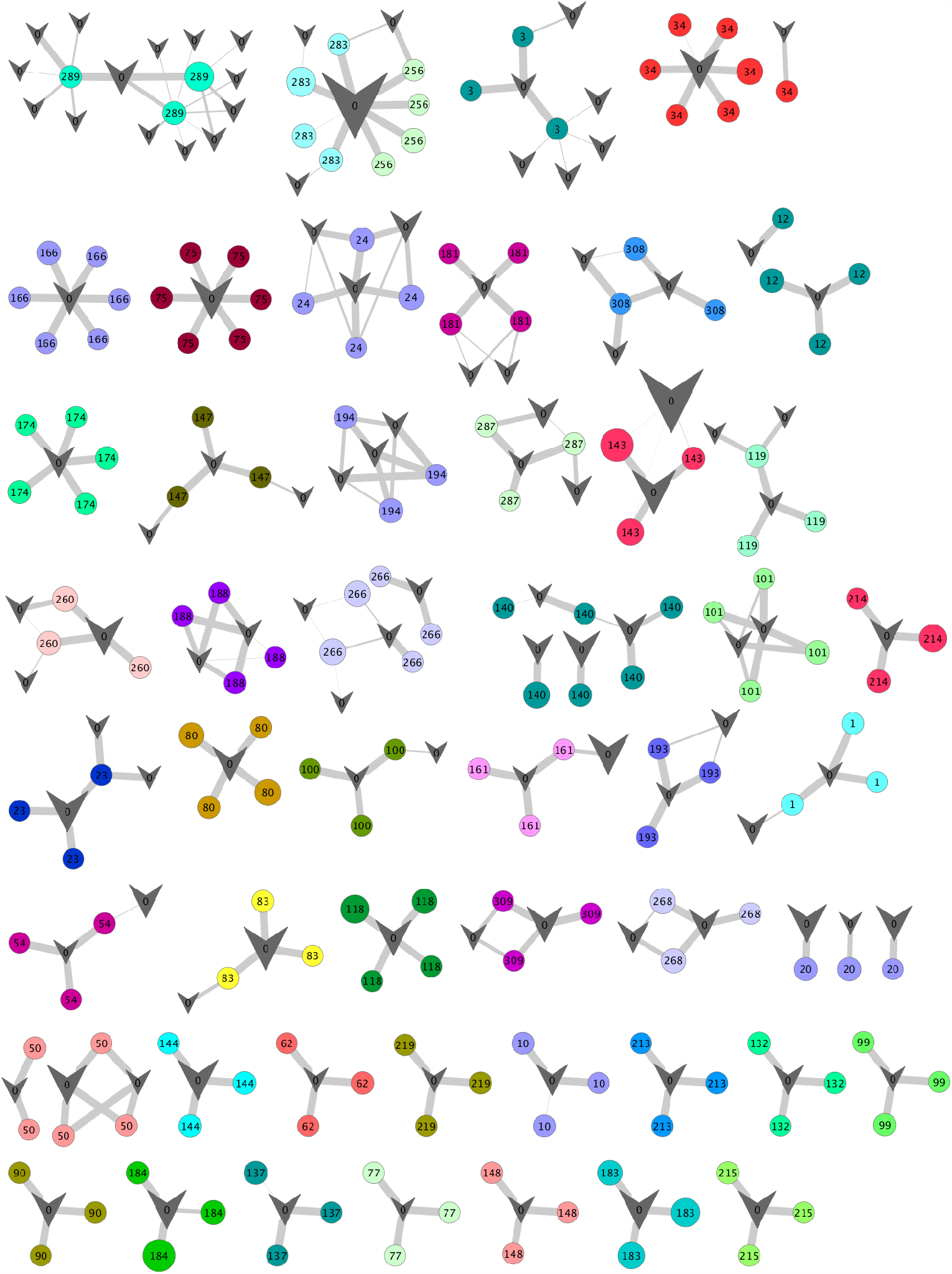
Viral MAG validation network. Network showing long-read validation of vMAG contig clusters from a sheep fecal metagenome. Viral contigs (circles, labeled with vMAG identifiers) are randomly colored according to the vMAG they belong to and linked to reference long-read viral genomes that they aligned to (grey check marks). The node size represents the viral sequence length, and the edge weight represents the percent of the short-read viral contig that aligned to the long-read reference. A random subset of 50 vMAGs was chosen for this visualization from a pool of vMAGs with at least 3 contigs and with at least one reference found for each contig.

**Fig. S4.**
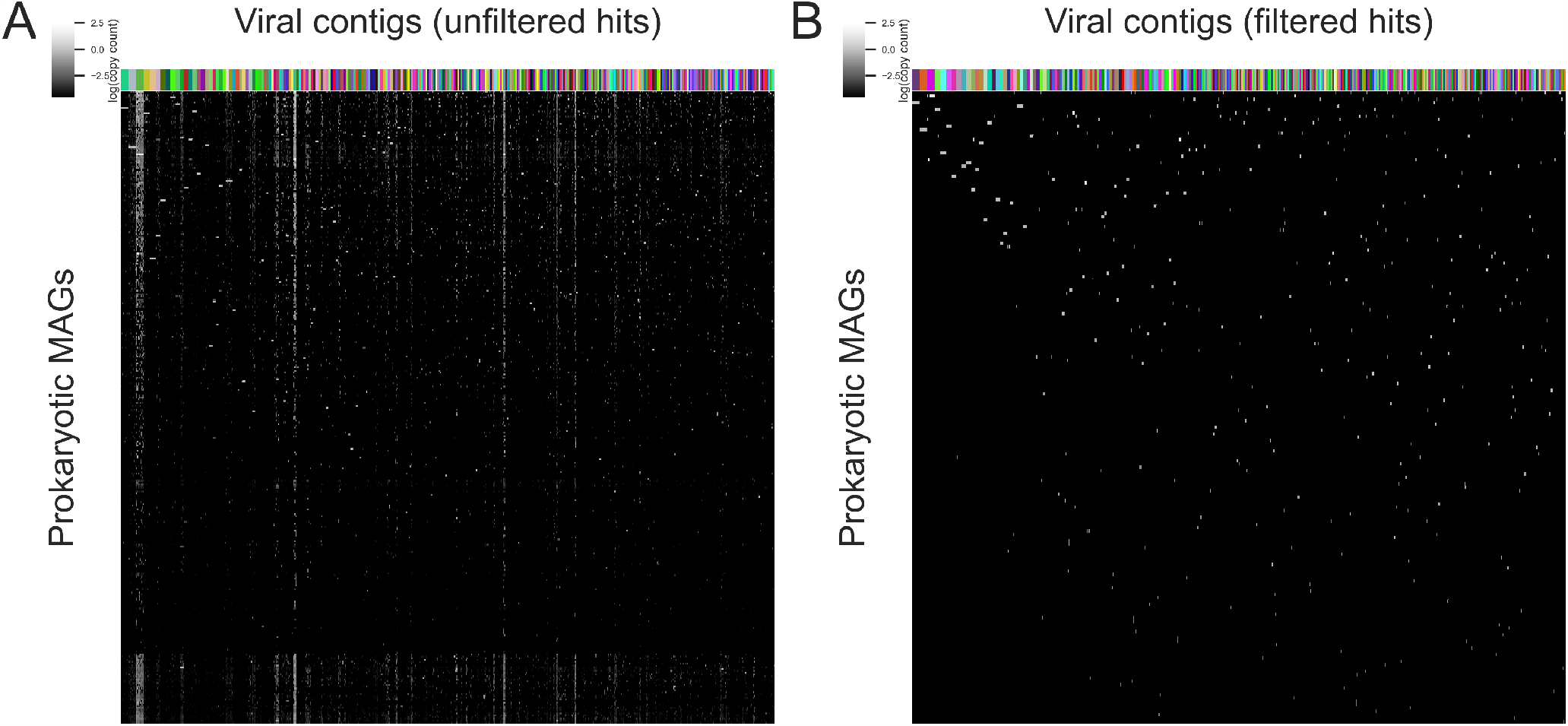
Viral contig host predictions. Prokaryotic hosts identified for viral contigs with ProxiPhage from a sheep fecal metagenome with ProxiPhage before (A) and after (B) thresholding. The color map encodes for the log of the estimated average copy count of each phage genome in its host. Columns are clustered according to vMAG membership (labeled with random colors) and rows are grouped based on linkage similarity with seaborn clustermap. Only viral contigs from viral MAGs are shown.

**Fig. S5.**
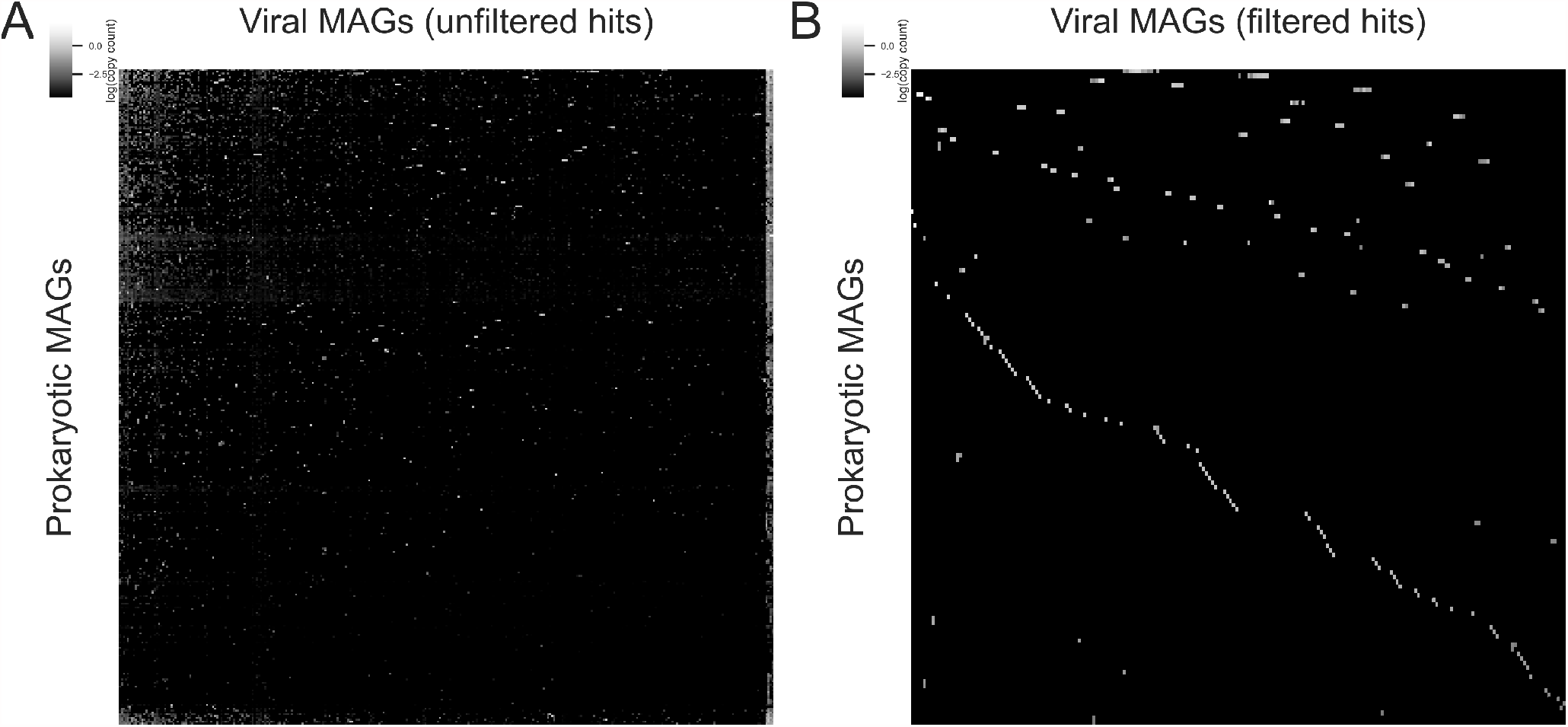
Viral MAG host predictions. Prokaryotic hosts identified for viral MAGs from a sheep fecal metagenome with ProxiPhage before (A) and after (B) thresholding. The color map represents the log of the estimated average copy count of each phage genome in its host. Rows and columns are clustered according to linkage similarity with seaborn clustermap.

